# PaIRKAT: A pathway integrated regression-based kernel association test with applications to metabolomics and COPD phenotypes

**DOI:** 10.1101/2021.04.23.440821

**Authors:** Charlie M. Carpenter, Weiming Zhang, Lucas Gillenwater, Cameron Severn, Tusharkanti Ghosh, Russel Bowler, Katerina Kechris, Debashis Ghosh

## Abstract

High-throughput data such as metabolomics, genomics, transcriptomics, and proteomics have become familiar data types within the “-omics” family. For this work, we focus on subsets that interact with one another and represent these “pathways” as graphs. Observed pathways often have disjoint components, i.e. nodes or sets of nodes (metabolites, etc.) not connected to any other within the pathway which notably lessens testing power. In this paper we propose the <u>Pa</u>thway <u>I</u>ntegrated <u>R</u>egression-based <u>K</u>ernel <u>A</u>ssociation <u>T</u>est (PaIRKAT), a new kernel machine regression method for incorporating known pathway information into the semi-parametric kernel regression framework. This paper also contributes an application of a graph kernel regularization method for overcoming disconnected pathways. By incorporating a regularized or “smoothed” graph into a score test, PaIRKAT is capable of providing more powerful tests for associations between biological pathways and phenotypes of interest and will be helpful in identifying novel pathways for targeted clinical research. We evaluate this method through several simulation studies and an application to real metabolomics data from the COPDGene study. Our simulation studies illustrate the robustness of this method to incorrect and incomplete pathway knowledge, and the real data analysis shows meaningful improvements of testing power in pathways. PaIRKAT was developed for application to metabolomic pathway data, but the techniques are easily generalizable to other data sources with a graph-like structure.

**Author Summary:** PaIRKAT is a tool for improving testing power on high dimensional data by including graph topography in the kernel machine regression setting. Studies on high dimensional data can struggle to include the complex relationships between variables. The semi-parametric kernel machine regression model is a powerful tool for capturing these types of relationships. They provide a framework for testing for relationships between outcomes of interest and high dimensional data such as metabolomic, genomic, or proteomic pathways. Our paper proposes PaIRKAT, a method for including known biological connections between high dimensional variables by representing them as edges of ‘graphs’ or ‘networks.’ It is common for nodes (e.g. metabolites) to be disconnected from all others within the graph, which leads to meaningful decreases in testing power whether or not the graph information is included. We include a graph regularization or ‘smoothing’ approach for managing this issue. We demonstrate the benefits of this approach through simulation studies and an application to the metabolomic data from the COPDGene study.

## Introduction

Metabolomics is the study of the metabolite composition of a cell, tissue, or biological fluid. Leading metabolomic experimental techniques such as liquid or gas chromatography coupled with mass spectrometry (LC-MS or GC-MS) and nuclear magnetic resonance (NMR) spectroscopy are able to capture the abundance of all metabolites within a cell (the metabolome). These technologies provide high-throughput data similar to other familiar -omics datatypes such as genomics, transcriptomics, and proteomics. An important advantage of metabolomics over other -omics data is its proximity to biological phenotypes(1). While genomic or proteomic data are vital pieces for understanding the progression from DNA to phenotype, the metabolites are the end products of the enzymatic reactions of a cell(2). The metabolome is comprised of exogenous (environmentally derived) and endogenous (genetically regulated) metabolites which can be used as biomarkers for the current phenotypic state of a cell or organism.

Like other -omics data, careful considerations of the metabolome’s unique characteristics are required to fully leverage it for biological insights. Specifically, metabolites are known to be related directly and indirectly by enzymatic reactions within a metabolomic pathway. These connections are sometimes handled separately through a secondary pathway enrichment analysis(3). Clustering methods have been developed to incorporate this connectivity into the primary analysis in order to avoid this two-step approach. These include Bayesian methods for metabolite clustering based on peak detection(4,5) and *ad hoc* methods based on singleton metabolite presence(6). For this work, we choose to group subsets of metabolites that interact with one another and represent these pathways as *graphs* or *networks*. Throughout this paper we will use the term *graph* and *network* interchangeably. Open source databases with metabolomic pathway documentation such as the Kyoto Encyclopedia of Genes and Genomes (KEGG), the Human Metabolome Database (HMDB), Reactome, OmniPath, and WikiPathways are growing resources(7–11), and the pathways within these databases are easily translated to graphs to be used in downstream analyses.

The semiparametric kernel machine regression method(12,13) has gained popularity in many areas of biomedical research such as genomics, microbiome analysis, and neuroimaging(14–16). One reason for its popularity is that it provides a computationally scalable method of classification and regression through the introduction of a *kernel* function. Another is that it provides a setting for formal statistical estimation and testing procedures for high-dimensional data sources, often through the use of a score statistic. Formal statistical tests are useful for metabolomic research, as a goal is often identifying specific metabolites and pathways for further inquiry. At a high level, kernel machines test for relationships between an outcome and a set of predictors by testing if variation between the two correspond with one another.

A hurdle more unique to metabolomics is the high levels of sparsity in individual metabolites and pathway connectivity. While metabolomic databases (e.g. KEGG, HMDB) are growing, none are considered complete. Data generating techniques like LC-MS and GC-MS are also imperfect technologies that may miss metabolite abundances that are too low(17). Thus, pathway representations of metabolomic data are often sparse and disconnected, i.e. nodes or sets of nodes (metabolites, etc.) not connected to any other within the pathway.

Disjoint nodes are of concern for graph-structured data since many techniques for inference on graph-structured data involve spectral decomposition and require full rank matrices from a fully connected graph. Techniques that force graphs to be fully connected by making small, uniform changes to the structure have been suggested for handling this issue(18,19). However, it is understood that these alterations impose new challenges by changing the subspaces spanned by the graph. Freytag et al. developed a network-based kernel for genomic pathway analysis in which they impose “as much noise as necessary” to ensure positive semidefinite matrices(20). This method and others like it are tailored to genomic data and not applicable to other omics data. The *PIMKL* method works with pathways within the metabolome by combining them through a weighted summed kernel(21). These weights provide insight into the importance of each sub-pathway, but this does not surmount to the level of evidence gathered from a direct comparison between specific pathways and phenotype.

In this paper we propose the <u>Pa</u>thway <u>I</u>ntegrated <u>R</u>egression-based <u>K</u>ernel <u>A</u>ssociation <u>T</u>est (PaIRKAT), a new kernel machine regression method for incorporating known pathway information into the semi-parametric kernel regression framework. In addition, PaIRKAT contributes an application of a graph kernel regularization method for overcoming sparse connectivity and disjoint pathways. To our knowledge, this is the first method to incorporate graph regularization into a kernel regression test. PaIRKAT allows for tests of association with phenotypes and the specific pathways while integrating pathway structure, and, instead of adding small amounts of noise, this approach dampens noisy components of a pathway while preserving biologically relevant signals. This leads to improved testing power and better overall biomarker detection. We evaluate these methods through several simulation studies and an application to real metabolomics data from the COPDGene(22) study.

## Results

### Method Overview

Here we provide the main steps of PaIRKAT and provide an overview of the ideas behind them. The method is described in full in **Methods and Models**. The primary goal of PaIRKAT is to include the topographical information of graph structured data into the kernel machine regression model. We use the semiparametric kernel machine model to test for relationships between the phenotype of interest, ***Y***, and high dimensional data, ***Z***, while controlling for important covariates, ***X***, in the model *g*(***Y***) = **Xβ** + *h*(***Z***) + ***ϵ***. In this model *h*(·) is a positive semidefinite kernel function that transforms ***Z*** to a feature space.

Omics data (metabolomics, genomics, etc.) can often be represented as a graph (or network) with edges representing biological interactions between the nodes (metabolites, etc.). The topography of a graph is often captured through its *normalized Laplacian*, 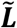. We explored the utility of incorporating the Laplacian directly into the kernel machine but found it to be ineffective. Instead, we transform 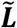 using methods designed to dampen noisy aspects of a graph while preserving its biologically relevant features. The PaIRKAT method is to include this *regularized Laplacian*, 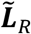, in the model through the kernel function thusly 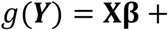 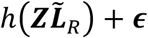. Tests for relationships between ***Y*** and 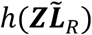 are accomplished using a score test based on a Satterthwaite approximation of a *χ*^2^ distribution.

### Simulation Results

A complete description of our simulation study can be found in **Methods and Models**, but we give a brief synopsis of the simulation scheme. We first randomly generated a graph. Second, we randomly generated features, ***Z***, from multivariate normal distribution with a covariance structure derived from the graph. Lastly, we randomly generated a normally distributed outcome, ***Y***, with a mean based on a linear relationship between the columns of ***Z***. We performed tests ignoring graph topography, including graph topography via the normalized Laplacian 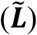, and our proposed method PaIRKAT of including graph topography via the regularized Laplacian 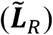. Our simulations aimed to assess how sensitive our method is to incomplete and/or incorrect graph information.

Type I error rates for PaIRKAT are summarized in Tables 1, 2, 3, and 4. The type I error rates for tests using a graph’s normalized Laplacian, 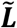 (see Methods section for definition), are summarized in Tables S1, S2, S3, and S4. The type I error rate of ≈ 0.05 is maintained throughout all simulation scenarios.

**Table 1:** Type 1 error rates using complete pathways. Error rates were calculated from score tests on 1000 simulated data sets. All simulations used graphs with 15, 30, or 45 nodes. No nodes or edges were dropped for these simulations.

|  | Pathway size |  |  |
| --- | --- | --- | --- |
|  | 15 | 30 | 45 |
| *Perfect* | 0.073 | 0.051 | 0.067 |
| *Partial 10%* | 0.051 | 0.049 | 0.059 |
| *Partial 40%* | 0.054 | 0.061 | 0.055 |
| *Partial 70%* | 0.054 | 0.061 | 0.055 |
| *Complete Mismatch* | 0.058 | 0.047 | 0.054 |
| *No Pathway* | 0.059 | 0.058 | 0.055 |

**Table 2:** Type 1 error rates using pathways with 5% missing edges. Error rates were calculated from score tests on 1000 simulated data sets using graphs with 15, 30, or 45 nodes. The graph used to simulate ***Z*** and ***Y*** was of medium edge density, while the graph used to test was of low density. The low-density graphs are drawn from the Barabasi-Albert model with edge density 0.13, 0.07, and 0.04 for graphs with 15, 30, and 45 nodes, respectively. <u>Medium edge</u> density graphs are created by giving any 2 nodes without a direct edge between them a 5% chance of becoming directly connected. This creates graphs with an average edge density of 0.18, 0.12, and 0.09 for graphs with 15, 30, and 45 nodes, respectively.

|  | Pathway size |  |  |
| --- | --- | --- | --- |
|  | 15 | 30 | 45 |
| *Perfect Network* | 0.052 | 0.056 | 0.046 |
| *Partial 10 Mismatch* | 0.046 | 0.063 | 0.047 |
| *Partial 40 Mismatch* | 0.051 | 0.068 | 0.052 |
| *Partial 70 Mismatch* | 0.051 | 0.068 | 0.052 |
| *Complete Mismatch* | 0.052 | 0.060 | 0.064 |
| *No Network* | 0.051 | 0.048 | 0.044 |

**Table 3:** Type 1 error rates using pathways with 15% missing edges. Error rates were calculated from score tests on 1000 simulated data sets using graphs with 15, 30, or 45 nodes. The graph used to simulate ***Z*** and ***Y*** was of high edge density, while the graph used to test was of low density. The low density graphs are drawn from the Barabasi-Albert model with edge density 0.13, 0.07, and 0.04 for graphs with 15, 30, and 45 nodes, respectively. <u>High edge</u> density graphs are created by giving any 2 nodes without a direct edge between them a 15% chance of becoming directly connected. This creates graphs with an average edge density of 0.26, 0.21, and 0.19 for graphs with 15, 30, and 45 nodes, respectively.

|  | Pathway size |  |  |
| --- | --- | --- | --- |
|  | 15 | 30 | 45 |
| *Perfect Network* | 0.061 | 0.059 | 0.064 |
| *Partial 10 Mismatch* | 0.039 | 0.057 | 0.059 |
| *Partial 40 Mismatch* | 0.045 | 0.065 | 0.058 |
| *Partial 70 Mismatch* | 0.045 | 0.065 | 0.058 |
| *Complete Mismatch* | 0.062 | 0.046 | 0.054 |
| *No Network* | 0.048 | 0.055 | 0.053 |

**Table 4:**
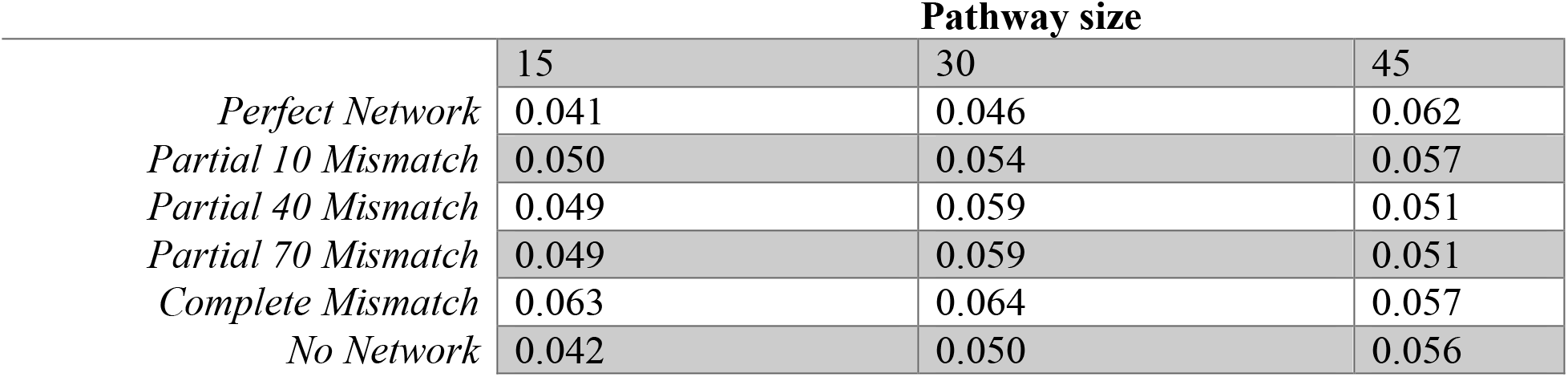
Type 1 error rates using pathways with dropped nodes. Error rates were calculated from score tests on 1000 simulated data sets using graphs 15, 30, or 45 nodes initially. The graph used to simulate ***Z*** and ***Y*** contained all nodes. Nodes with degree below the 25^th^ percentile within a graph had a 25% chance of being dropped before testing.

The power curves for all pathway structures while simulating complete knowledge, missing edges, and missing nodes are displayed in Figure 1. Having a perfect pathway structure provides the most power. Relationships between an outcome and pathway are easier to detect in larger pathways. The more incorrect direct edges in the pathway, the lower the overall power. We also see that increasing the overall signal to noise ratio improves power for all pathway structures (Figure 2). PaIRKAT 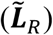 achieves approximately 80% power at a signal to noise ratio around 0.23, whereas ignoring network information requires a signal to noise ratio nearly three times large, about 0.60, and only including the Laplacian never achieves 80% power (Figure 2).

**Fig 1:**
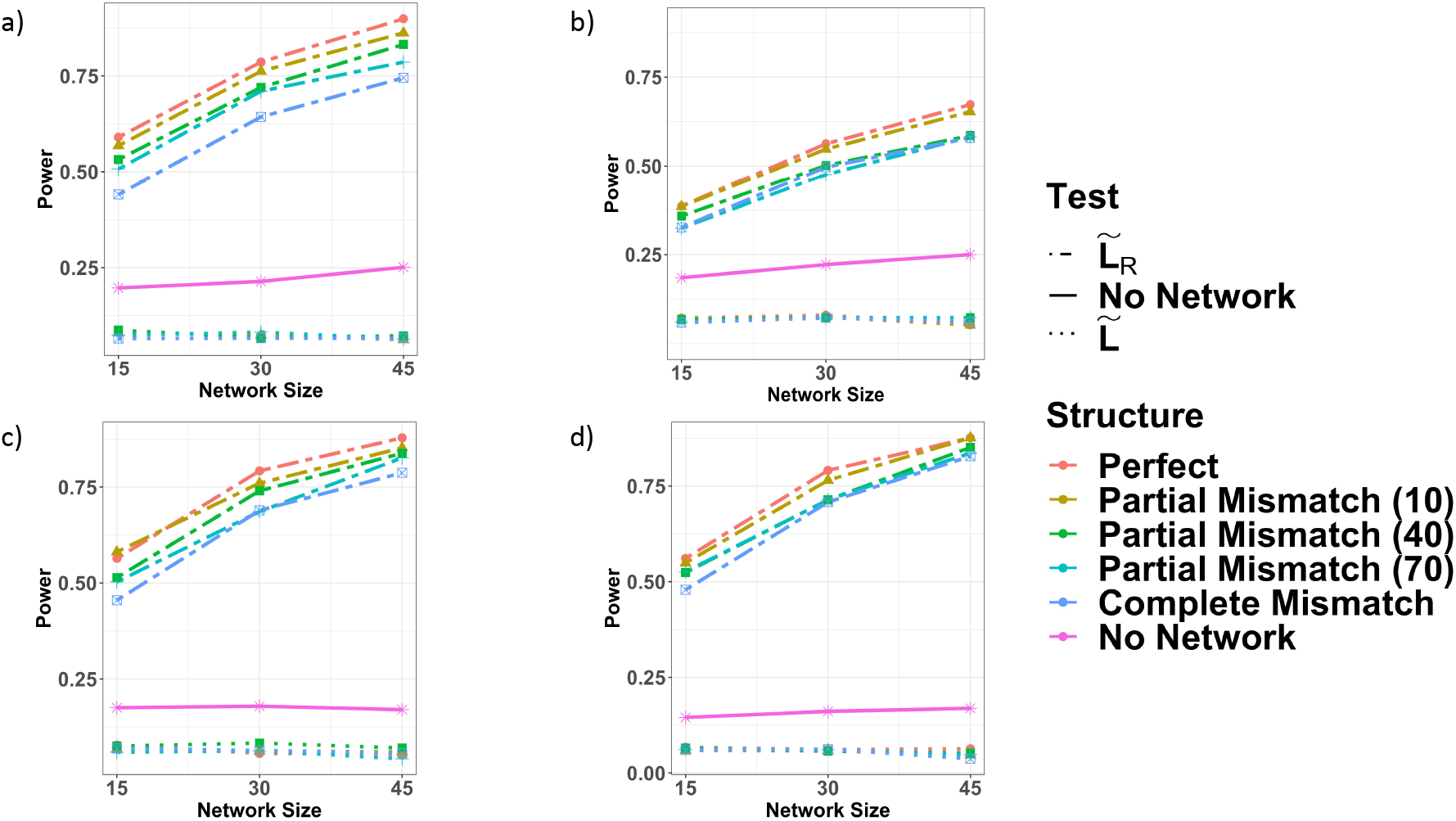
Power curves from the four pathway *knowledge* and 6 pathway *structure* simulation scenarios. Power curves were all calculated from score tests on 1000 simulated data sets using graphs with 15, 30, or 45 nodes. Power curves assuming complete pathway knowledge with no dropped edges or nodes are displayed in a). For (b) and (c), the graph used to simulate ***Z*** and ***Y*** was of medium or high density, respectively, while the graph used to test was of low density. Medium and high edge density graphs used for data generation had ~5% and ~15% more edges, respectively, than the low density graph used for testing. The power curve generated assuming missing nodes (d) used all graph nodes to generate ***Z*** and ***Y***. Then nodes (and corresponding columns of ***Z***) with degree below the 25^th^ percentile within a graph had a 25% chance of being dropped before testing.

**Fig 2:**
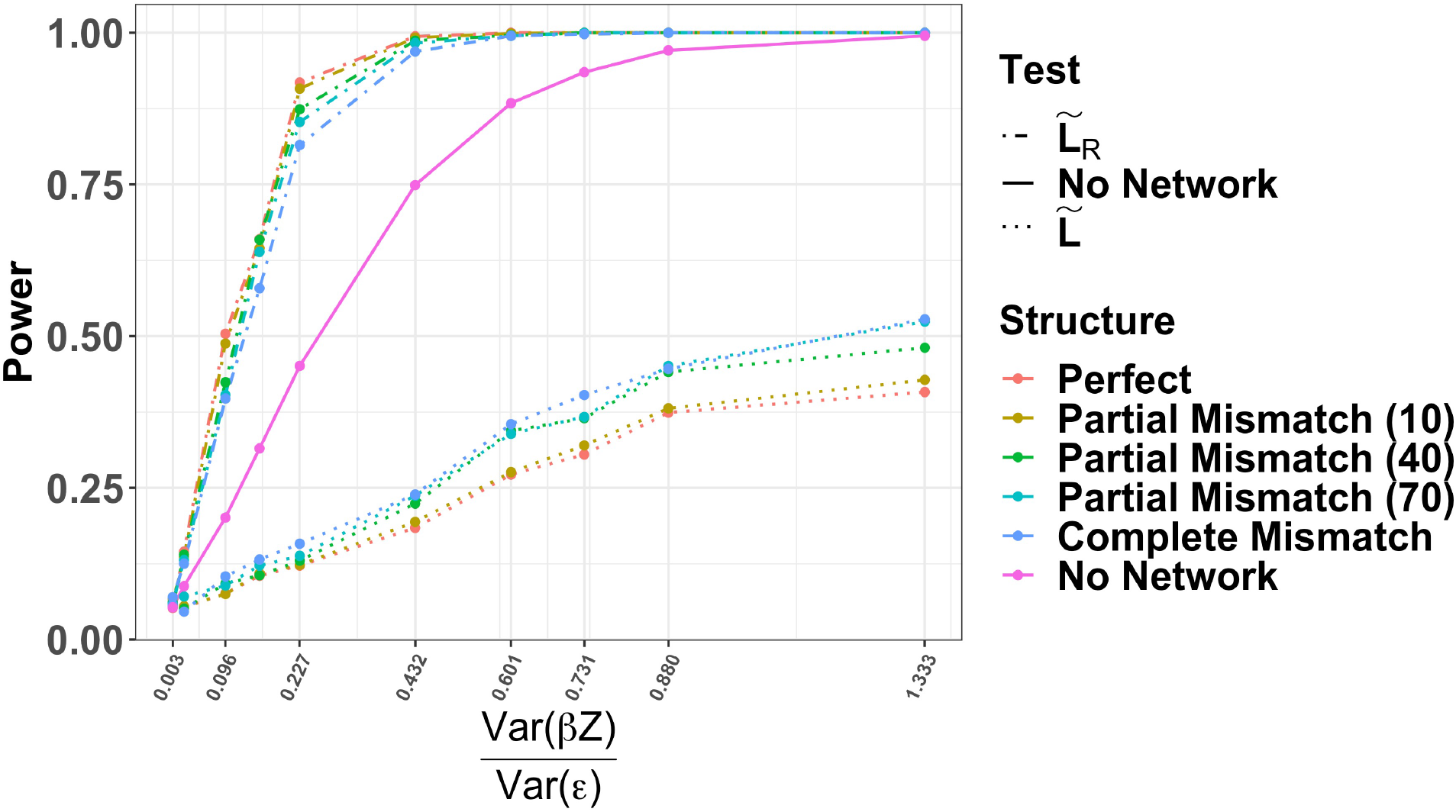
Signal to Noise Ratio. Power curves from increasing the signal to noise ratio while assuming complete pathway knowledge. The signal to noise ratio was calculated as the as the ratio between the overall variance in ***Y***, 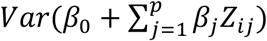, and the overall residual variance, 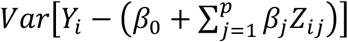. Each power calculation comes from score tests on 1000 simulated data sets using graphs with 30 nodes.

### COPDGene Analysis Results

A complete description of these analyses can be found in **Methods and Models**, but here we give a brief description of the outcome variables we analyzed. We create models for two phenotypes from the COPDGene study(22): (1) percent emphysema and (2) the ratio of post-bronchodilator forced expiratory volume at one second divided by forced vital capacity (FEV_1_/FVC). To normalize FEV_1_/FVC, we use the following log ratio transformation, 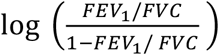. This is referred to as the “log FEV_1_/FVC ratio” for simplicity. We test for associations between 28 pathways and each outcome under the same three conditions in the simulation study: ignoring graph topography, including graph topography via the normalized Laplacian 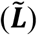, and our proposed method PaIRKAT of including graph topography via the regularized Laplacian 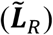.

Including the metabolites’ regularized graphs had large impacts on the associations between the log FEV_1_/FVC ratio and several subsets of metabolites. For the 28 pathways tested, power was improved for 18 pathways when using PaIRKAT vs. using 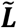 or ignoring pathway information. Of note, the strength of the associations between the log FEV_1_/FVC ratio and the *ABC transporters,* the *arginine and proline metabolism,* the *cysteine and methionine metabolism,* the *pyrimidine metabolism,* the *glycine, serine, and threonine metabolism,* and the *neuroactive ligand-receptor interaction* metabolite subsets increased dramatically. The average value was also lower for 5 pathways with using 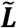 vs. ignoring pathway information. However, there was not a significant result from any method for 4 of these pathways, and PaIRKAT provided similar power for the fifth. Figure S1 displays the p-values from the kernel regression tests for associations between the log FEV_1_/FVC and the 28 pathways of interest for each subsample size.

Including the metabolites’ regularized graphs also had impacts on the associations between percent emphysema and several subsets of metabolites. For the 28 subsets of metabolites tested, power was improved for 17 pathways when using PaIRKAT vs. including 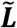 or ignoring pathway information. Of note, the strength of the associations between percent emphysema and the *ABC transporters, the B-alanine metabolism, the neuroactive ligand248 receptor interaction, the glycine, serine and threonine metabolism, and the histidine metabolism* metabolite subsets increased dramatically when using PaIRKAT vs. ignoring pathway information. The average p-value was again lower for the same 5 pathways with using 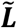 vs. ignoring pathway information. However, there was still not a significant result from any method for 4 of these pathways, and PaIRKAT provided similar power for the fifth. Figure S2 displays the p-values from the kernel regression tests for associations between percent emphysema and the 28 pathways of interest for each subsample size.

Figure 3 displays results from 3 pathways selected to illustrate PaIRKAT’s impact on power for fully connected (left column), partially disconnected (middle column), and sparse (right column) graphs. For the *steroid hormone biosynthesis* pathway, an almost completely sparse pathway, we see virtually no differences between PaIRKAT and ignoring pathway connectivity. We also see relatively small differences between all three methods for the fully connected *aminoacyl-tRNA biosynthesis* pathway. The major impacts from PaIRKAT come when there are a few nodes or node subsets disjoint from the rest of the graph, as we see in the *cysteine and methionine metabolism*.

**Figure 3:**
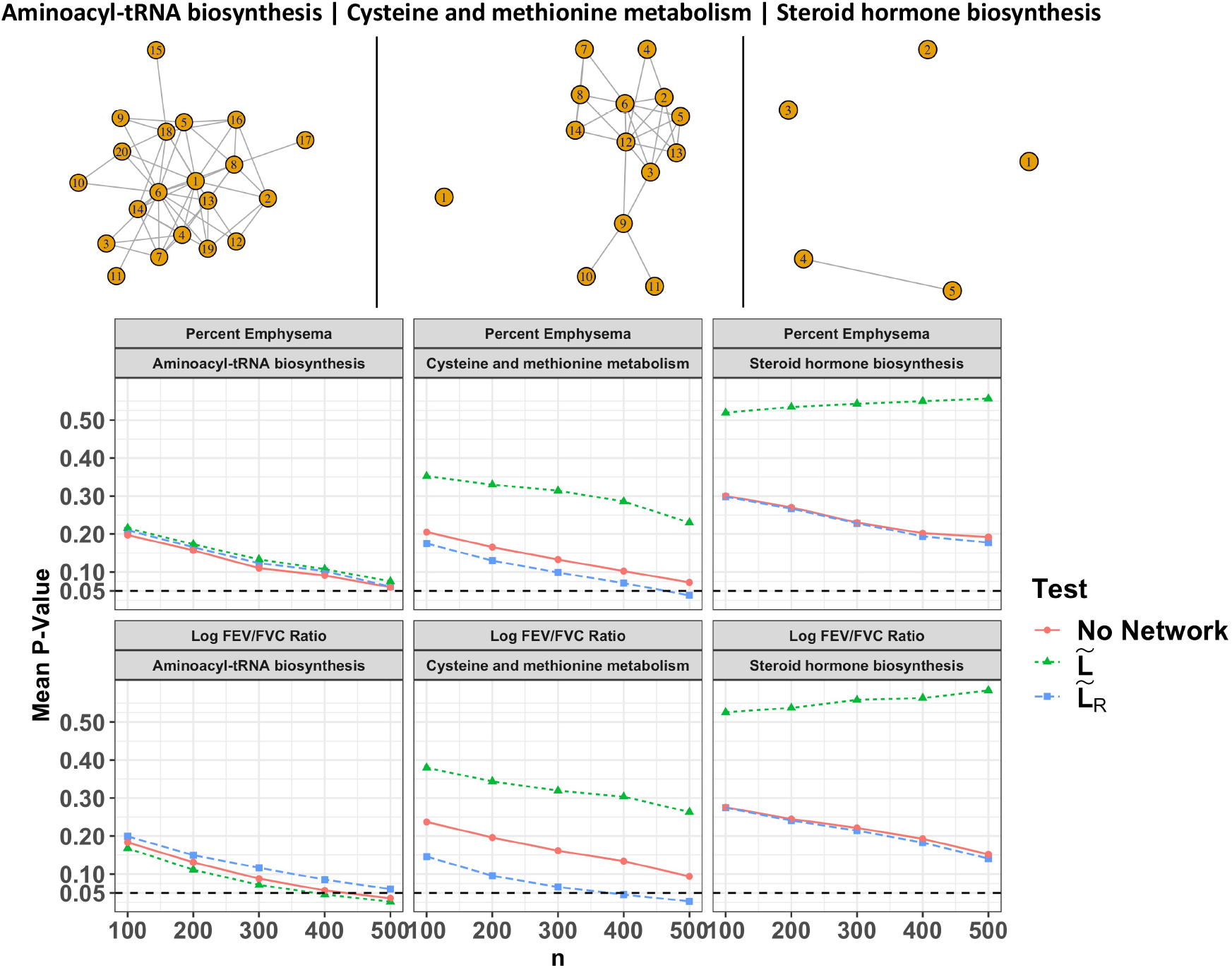
Selected results from COPDGene subset analysis. Average p-values from kernel regressing tests that do not include pathway information (No Laplacian, red circles), include pathway information through a normalized Laplacian (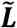, green triangles), and include pathway information through a regularized normalized Laplacian (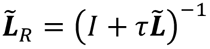, blue squares) are displayed. P-values were averaged over 100 random subsets of size 100, 200, 300, 400, and 500 from the COPDGene dataset. *τ* was set to 1 for all tests that used 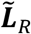. The 3 pathways selected illustrate the expected results in fully connected (left), partially disconnected (middle), and sparse (right) graphs.

## Discussion

We developed PaIRKAT, a method for incorporating pathway information under a kernel regression framework. Other methods to incorporate pathway connectivity via graph operations have been developed(20,21,23–26). PaIRKAT enables the researcher to test on specified pathways instead of aggregating all pathways through a weighted kernel as in(21,24). It can also handle disjointed pathways without adding in artificial noise to the network as in(18–20). This allows the investigator to compile information from multiple sources, e.g. KEGG and HMDB. The regression framework also expands upon a method developed for classification(23). Finally, this method provides a one-step testing procedure to simplify workflow compared to the popular two-step enrichment analysis procedure.

Pathway misspecifications from incomplete data collection or imperfect canonical pathways within databases are common hurdles in -omics studies. We explored the sensitivity of the method by simulating data assuming incorrect pathway structures and incomplete pathway knowledge. These studies show that our method is highly robust to pathway misspecifications. In smaller pathways, we see that the partially mismatched structure with ~10% of direct edges being incorrect does as well as the perfect network structure. This is likely due to the very small change from the perfect structure in these cases, as a graph with only 15 nodes could easily be unchanged with only a 10% chance to change an edge. Furthermore, even with incorrect or incomplete pathway information, our method provides significantly improved power over ignoring pathway information while maintaining an appropriate type I error rate. We believe this is because many indirect connections between nodes are preserved, and these connections still provide more accurate information than incorrectly assuming independence among nodes.

One benefit of using PaIRKAT is improved power to identify pathways that are associated with clinical phenotypes. For example, an application to the COPDGene dataset using KEGG’s database of metabolic pathways also illustrated PaIRKAT’s ability to improve testing power over simply treating metabolites as independent (Figure 3, S1, S2). The regularization technique was also able to handle pathways with few metabolites and/or disjoint components. Several tests had a dramatic boost in power from including pathway connectivity for both percent emphysema and the log FEV_1_/FVC ratio, and most pathways have been previously associated with COPD and lung function. Huang et al. linked environmental exposures, COPD risk, and metabolomic pathways, and found associations between COPD and the *histidine metabolism*, *cysteine and methionine metabolism*, and *β*-*alanine metabolism* pathways(27). The *glycine, serine and threonine metabolism*, *aminoacyl-tRNA biosynthesis*, *pyrimidine metabolism*, *pantothenate and CoA biosynthesis*, pathways have all previously been associated with asthma(28). The *β*-Alanine metabolism, ABC transporters, purine metabolism, pantothenate and CoA biosynthesis pathways were all differentially associated with COPD subclasses for patients with lung cancer(29). Another study of the COPDGene dataset(22) using a two-step pathway enrichment approach found that the *purine metabolism, mineral absorption, arginine biosynthesis, aminoacyl-tRNA biosynthesis, ABC transporters* and *glycine, serine and threonine metabolism* pathways were all associated with various measures of lung function and increased COPD exacerbations(30). The three *ABC transporters* have also been shown to be related to COPD in several murine knockout and human studies (see Chai et al.(31)). Finally, the *arginine biosynthesis* pathway has also been associated with COPD in multiple studies (32,33).

### Impacts of Regularization

In simulation studies and real data analyses we saw meaningful improvements in power by including pathway information through a graph’s regularized normalized Laplacian, (PaIRKAT) when compared to ignoring the pathway information or using 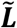. PaIRKAT was essential to maintaining testing power when graphs had disjoint nodes or sub-graphs. Using the normalized Laplacian, 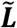, hindered testing performance compared to using PaIRKAT or ignoring the pathway information when a graph was disconnected. In connected graphs PaIRKAT, using 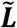, and ignoring the pathway information all performed similarly in the real data analyses (Figure 3).

It is well established that 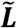 is a symmetric and positive semidefinite matrix with eigen values 0 ≤ *λ*_1_, *λ*_2_, … . *λ*_*p*_ ≤ 2, where the number of *λ*_*i*_ = 0 is the number of disjoint components of the undirected graph *G* (see **Methods and Models**). Therefore, graphs with very low connectivity, meaning many *λ*_*i*_ = 0, will not be as impacted by regularization since all *r*^−1^(*λ*_*i*_ = 0) = *a* for some scalar *a*. In words, there is no extra information from a graph when most nodes are disconnected from one another (e.g., Figure 3, right column).

In summary, our proposed method serves as a framework for including pathway information into a kernel machine regression test. We developed this method for application to metabolomic pathway data, but the techniques are easily generalizable to other data sources with a graph-like structure. It is important to examine the structure of a graph before applying a regularization step. Unique challenges arose from the sparsity present in many metabolomic pathways which can greatly hinder performance. We implement a graph regularization kernel to handle disconnected pathways. This regularization step is novel in the application of graph-based kernel machine regression to biological data. Our simulation studies illustrate the robustness of this method to improper and incomplete pathway knowledge. The method presented is capable of providing powerful tests for associations between biological pathways and phenotypes of interest and will be helpful in identifying novel pathways for targeted clinical research. The R code for simulations and an example workflow with source code is available at https://github.com/CharlieCarpenter/PaIRKAT, and an R shiny app is currently hosted at https://csevern.shinyapps.io/pairkat/.

## Methods and Models

### The Kernel Machine Model

We will assume that the data are properly filtered, imputed, and normalized for the methods described in this paper. Consider a dataset with observations from *n* subjects. Let ***Y*** be an *n* × 1 vector representing a continuous or discrete phenotype of interest. Also let ***X*** be a *n* × *q* matrix of clinical covariates and ***Z*** be an *n* × *p* matrix of metabolite abundances. The phenotype can then be modeled through the following semiparametric model

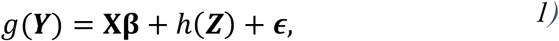

where *g* is any standard link function, **β** is a *q* × 1 vector of regression coefficients, ***ϵ*** is an *n* × 1 vector of normally distributed error terms, and *h* is a kernel function. There are no parametric assumptions placed on *h* except that it lies in some feature space. This more relaxed requirement from the kernel regression provides flexibility and robustness to model misspecification. Another key advantage of introducing the kernel function is its ability to capture nonlinear relationships between the phenotype (***Y*)** and the metabolome (***Z***) in a computationally tractable manner. These relationships are assumed to exist in some feature space, denoted 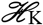, that is generated by a positive definite kernel function *K*(·,·). Under this condition, the representer theorem then allows *h*(***Z***) to be represented through the kernel function *K*(·,·) as 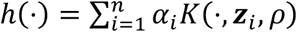 for some coefficients *α*_*i*_ ∈ ℝ. More detailed derivations can be found in texts by Schölkopf and Smola(34) as well as Cristianini and Shawe-Taylor(35).

The kernel function *K* can be thought of as a measurement of similarity of the metabolome between two individuals. Consequently, the choice of a kernel function defines the notion of similarity and covariance between the metabolome and the phenotype of interest. Two common choices for kernel functions are the *dth Polynomial Kernel*: 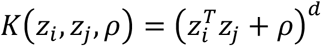 and the *Gaussian Kernel*: 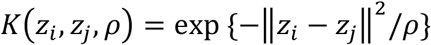, where ∥·∥ is the Euclidean (*L*_2_) norm. Both kernels contain a tuning parameter *ρ*. It is common in the machine learning literature for this parameter to be fixed to some ad-hoc value. For this work, we employ the Gaussian kernel and use the median of all pairwise Euclidean distances between metabolites as an empirical estimate of *ρ*. We choose to work with the Gaussian kernel since it is a *characteristic* kernel, a desirable property meaning that probability measures embedded onto 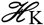 through the kernel function are unique. These familiar choices are used to illustrate our new method, but, in principle, several kernel functions and choices for estimating *ρ* could be applied.

### Kernel-based Score Test

Liu et al. show a connection between kernel machine regression and linear mixed models for semiparametric modeling of high dimensional data(12,13). The parameters ***β*** and *h*(***Z***) can be estimated by maximizing the scale penalized likelihood

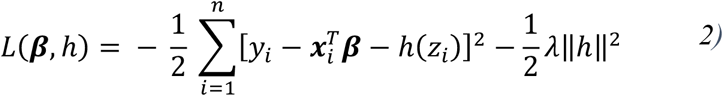

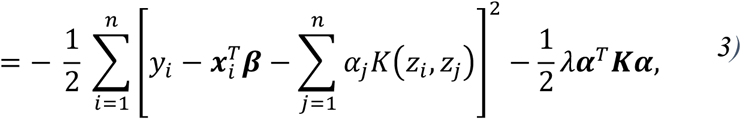

where ***K*** = *K*(*Z*_*i*_, *Z*_*j*_, *ρ*) is the semi-positive definite kernel function of choice. *h*(***Z***) can then be understood as subject specific random effects with mean 0 and variance *τ****K***. Testing for an association between phenotype and pathway is then equivalent to testing the null hypothesis *H*_0_: *τ* = 0 vs *H*_1_: *τ* > 0. Liu et al follow Zhang and Lin in the formulation of a score statistic for this variance component test(12,36). Specifically, the score statistic for *τ* is 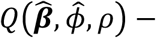 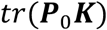 where

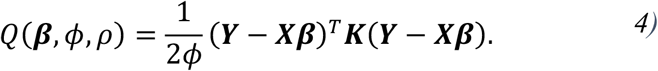

Since *Q* is a quadratic function of the residuals, an approximation for the null distribution can be obtained using Satterthwaite’s method of matching the mean and variance of *Q* and a scaled chi-squared distribution, 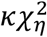. This results in 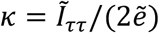 and 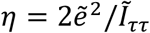 where 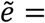 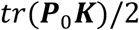 and 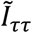 is the efficient information 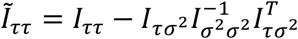. Here 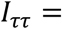 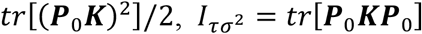, and 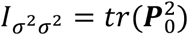. *ϕ* serves as a scaling parameter and ***P***_0_ is the projection matrix under the null model: *g*(***Y***) = **Xβ** + ***ϵ***. For continuous outcomes the projection matrix is ***P***_**0**_ = ***I*** − ***X***(***X***^***T***^***X***)^**−1**^***X***^**T**^ and *ϕ* is the residual variance, *σ*^2^. For dichotomous outcomes the projection matrix is ***P***_**0**_ = ***D***_**0**_ − ***D***_**0**_***X***(***X***^***T***^***D***_**0**_***X***)^**−1**^***X***^***T***^***D***_**0**_ and *ϕ* = 1, with 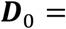 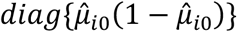 and 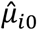 being the fitted probability *P*(*y*_*i*_ = 1) under the null model. A key advantage of working with the score statistic is the need to only fit the null model for testing, meaning 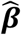 and 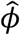 are their respective maximum likelihood estimates under the null model. Liu et al also demonstrated that this method is robust to the choice of *ρ*(12).

### Graph Laplacian

A network or graph, *G* = {*V*, *E*}, is a mathematical representation of any interconnected structure through a set *V* of *p* nodes (or vertices) and a set *E* of edges, where the elements of *E* are pairs 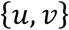 of distinct vertices, *u*, 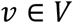. When applied to metabolomic pathways, nodes represent individual metabolites within the pathway and edges represent direct biochemical interactions/reactions between metabolites.

Two important features of any graph are its adjacency matrix, ***A***, and degree matrix, ***D***. ***A*** is a *p* × *p* matrix that is non-zero when an edge exists between two vertices. ***D*** is a *p* × *p* diagonal matrix with ***D***_[*i*,*i*]_ representing the number of nodes connected to node *i*. For this work, we represent pathways using undirected unweighted graphs, i.e. there is no ordering to the vertices defining an edge and 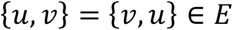. This means ***A*** will be a symmetric matrix with all entries either 1 or 0. Using these features, we can calculate a graph’s *Laplacian* ***L*** ≔ ***D*** − ***A*** and its *normalized Laplacian* 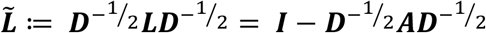, where ***I*** is a *p* × *p* identity matrix.

Both ***L*** and 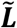 can be regarded as linear operators of functions ***f***: *V* → ℝ that induce a semi-norm ∥***f***∥_*L*_ = ⟨***f***, ***Lf***⟩ = ***f***^***T***^***Lf***. This semi-norm can be interpreted as a measure of “smoothness” or how much ***f*** varies over its domain. Standardizing ***L*** by the number of connections per node to obtain 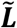 is a common approach in graph theory since 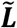 has several well-known and desirable properties. In particular, 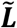 is symmetric and positive semidefinite, and its eigenvalues, *λ*_*i*_, are bounded such that they satisfy 0 ≤ *λ*_*i*_ ≤ 2 for *i* ∈ 1,2, … *p*(37).

### Graph Regularization

A key component of PaIRKAT is the ability to handle disjoint components within a graph. Pathway databases may not be complete, and untargeted data generating techniques may not be able capture all components within a pathway. This leaves some pathways with low connectivity and others with completely disconnected nodes. An interesting feature of a graph’s normalized Laplacian, 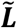, is that the number of disjoint pieces within a graph is captured by the number of 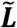 eigen values equal to 0. When *G* has disjoint nodes and disjoint nodes 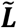 becomes less than full rank. This can lead to a decrease in our power to detect associations between phenotypes and metabolomic pathways. One proposed solution is to simply manipulate the adjacency matrix by adding a small constant to all entries(18,19), i.e. working with a modified adjacency matrix 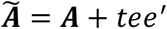, where *t* is a nonnegative tuning parameter and *e* is a vector of 1s. This yields a full rank matrix as desired, but we know that the subspace spanned by 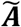 is not the correct subspace on which our graph lies.

A more elegant solution can be drawn from Smola and Kondor’s work on regularization of graphs(38) in which they draw on parallels between the standard Laplacian operator 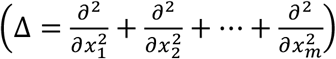 and the graph Laplacian to design regularization kernels for graphs. Rapaport, et al.(23) took a similar approach to graph smoothing, though this work was done in the context of classification not hypothesis testing. These ideas can be generalized further to represent any metric on a space. That is, for any two observations *i* and *j*, the inner product can be expressed as 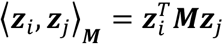, where ***M*** defines the metric on the vector space based on ∥***Z***_***i***_ − ***Z***_***j***_∥_***M***_. Purdom(39) presents this argument in the context of a “generalized” principal component analysis using a general metric ***M***. This can be seen as an application of a linear kernel on any metric space, whereas we apply the Gaussian kernels for hypothesis testing and, like Rapaport, focus on graph Laplacians for our metric.

For this work, we apply a regularization kernel to obtain a *regularized* normalized Laplacian: 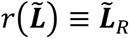. Regularizations of the Laplacian can be seen as regularizations of the eigenvalues of 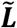, *r*(*λ*). There are many possible choices for *r*; the only requirement is that *r*^−1^(*λ*) > 0 for *λ* ∈ [0,2] to ensure 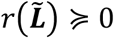. In classical Fourier analysis the size of *λ*_*i*_ ∈ [0,2] is directly proportional to the frequency of component *i* within Fourier space, which translates to the degree of noise within the system. This intuition tells us to limit *r*^−1^(*λ*) to monotonically increasing functions in order to impose higher penalties to more uneven portions of the graph while preserving the lower frequency components, which we assume translate to the prevalent biological signals. Smola and Kondor recommend further limiting choices of *r* to functions expressible by power series such as a *diffusion* kernel(40), 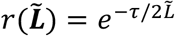. See their paper for complete details on the derivation of different regularization functions.

PaIRKAT implements a “linear” regularization function

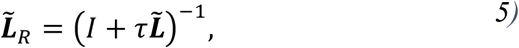

where *τ* > 0 is a bandwidth parameter and *I* is a *p* × *p* identity matrix. We choose this regularization for its simplicity and interpretability of *τ*. Increasing *τ* linearly increases the amount of smoothing performed in *r*^−1^(*λ*) = 1 + *τλ*. With the graph’s sparse connectivity accounted for through the linear regularization function, we can now conduct a kernel machine test while incorporating connectivity within a pathway through 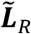 into (1) thusly

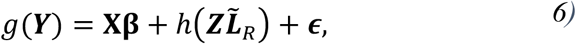

where *h* is a kernel function applied to 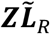 and the other model components are as described in (1). The kernel-based score test can then be applied to obtain powerful tests for associations between connected or disconnected pathways and a phenotype of interest.

### Simulation Study

#### Simulation Scenarios

We conducted multiple simulation studies to assess whether or not the proposed method is robust to imperfect pathway information. We assumed 3 different “pathway knowledge” scenarios and 4 different “pathway structure” scenarios (Figure 4). Different pathway knowledge scenarios refer to different types of missing information, whereas pathway structure scenarios refer to different configurations of the “known” nodes and edges. We simulate using both the normalized Laplacian, 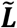, and PaIRKAT’s regularized normalized Laplacian, 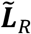, as well as ignoring the pathway information.

**Fig 4:**
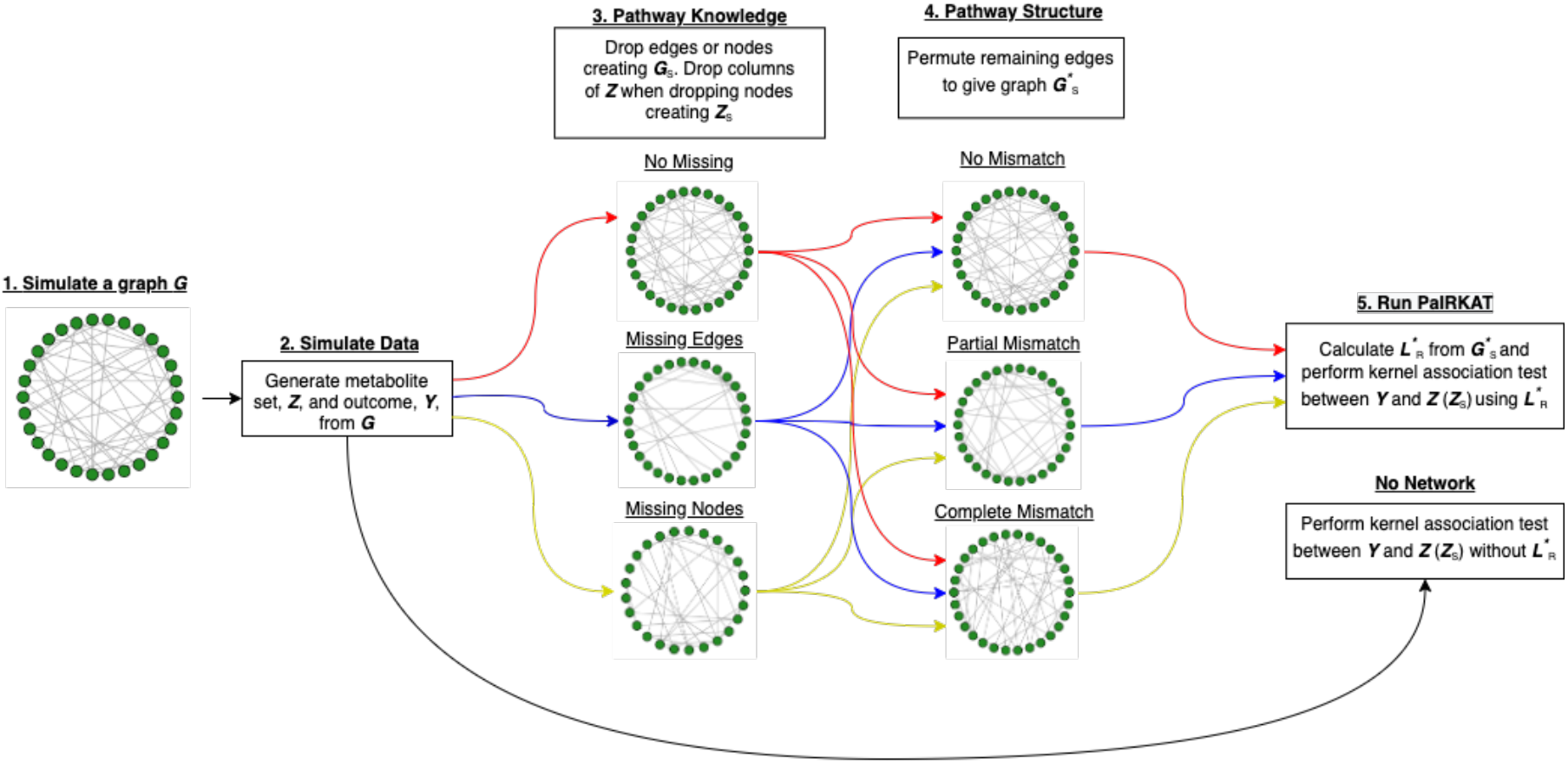
Flowchart of simulation procedure. We (1) simulate a graph *G*, (2) generate ***Z*** and ***Y*** from *G*, (3) drop nodes or edges from *G* to give a smaller graph *G*_*s*_ (drop corresponding columns of ***Z*** when dropping edges to create ***Z***_*s*_), (4) permute edges to create an improperly structured graph 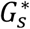, (5) calculate the regularized normalized Laplacian 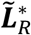 from 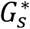, and finally (5) test for an association between 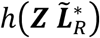 (or 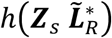) and ***Y*** in the model 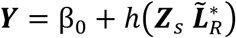. For the “no network” simulations, we only use step (1), step (2) and step (5) without including 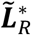.

#### Pathway Knowledge

We simulated three different *knowledge* scenarios to represent incomplete pathway database information and/or incomplete data collection.

1. <u>No missing</u>: Assuming the nodes measured (metabolites, genes, etc.) and edges connecting them are a perfect representation of the biological pathway of interest.
2. <u>Missing edges</u>: Assuming that some biological interactions (edges) are missing from the documented pathway. Here we generate a graph *G* = {*V*, *E*} according to the Barabasi-Albert model for a “low” edge density. We then give every set 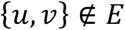 a 5% or 15% percent chance of being added to *E* for a “medium” or “high” edge density graph, respectively. ***Z*** and ***Y*** are then generated from the medium or high edge density graph, but 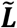 or 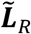 is calculated from the original “low” edge density graph. Examples of these graphs are shown in Figure S3.
3. <u>Missing nodes</u>: Assuming that some of the nodes (and hence their edges) are missing from the documented pathway. Here a graph is used to generate ***Z*** and ***Y***. Then nodes with degree below the 25^th^ percentile have a 25% chance of being removed before calculating 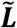 or 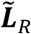. The corresponding columns and rows of ***Z*** and 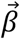 are removed as well. Examples of these graphs are shown in Figure S4.

#### Pathway Structures

After we simulate a pathway knowledge scenario, we alter the pathway *structure* to represent incorrect edge connections within a database. Examples of structures 1, 2, and 3 are displayed in Figure 5.

1. <u>No mismatch</u>: No alterations to graph edges. The graph used to simulate ***Z*** and ***Y*** is the same graph used to calculate 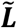 or 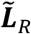 (Figure 5, left).
2. <u>Partial Mismatch</u>: a graph, *G*_1_ = {*V*_1_, *E*_1_}, is used to simulate ***Z*** and ***Y***. This graph’s edges are permuted such that any edge 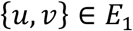 has a 10%, 40%, or 70% chance of being changed to some {*u, W*} ∉ *E*_1_; i.e. approximately 10%, 40%, or 70% of direct edges are incorrect before calculating 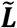 or 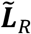 (Figure 5, middle).
3. <u>Complete Mismatch</u>: a network *G*_1_ = {*V*_1_, *E*_1_} is used to simulate ***Z*** and ***Y***. A new random graph, *G*_2_, is then draw and forced to have no edges that match *G*_1_, i.e. *V*_1_ = *V*_2_ but if 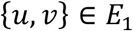 then 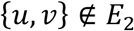. We then calculate 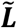 or 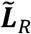 from *G*_2_ (Figure 5, right).
4. <u>No Pathway</u>: a graph is used to simulate ***Z*** and ***Y***. This connectivity is ignored while testing by not including 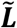 or 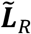 in the kernel function.

**Fig 5:**
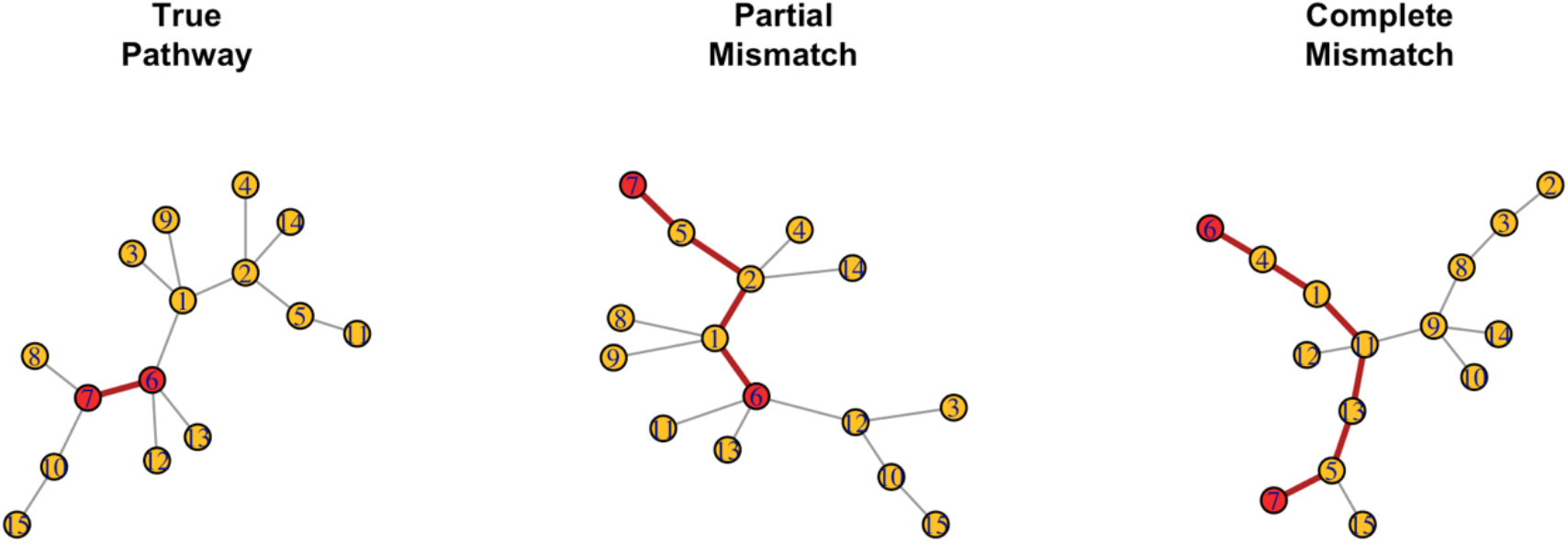
Examples of the three different pathway *structures*. Nodes 6 and 7 are highlighted in red to help display the effects of different pathway structures. (Left) The “true” pathway or graph that is used to simulated ***Z*** and ***Y***. This is the graph used for tests under a “perfect pathway structure” scenario. (Middle) A graph with approximately 40% of the edges from the “true” graph directly connecting the wrong nodes. This is used for tests under a “partial mismatch (40) structure” scenario. (Right) A graph with 0 shared edges with the “true” graph. This is the graph used for tests under a “complete mismatch structure” scenario.

All pathway *structures* were considered under each different pathway *knowledge* scenario. The different pathway structures were imposed after simulating under different pathway knowledge assumptions. Each simulated pathway structure and knowledge combination followed 5 steps: (1) simulate a graph *G*, (2) generate ***Z*** and ***Y*** from *G*, (3) drop nodes and/or edges (based on *knowledge* assumption) from *G* and ***Z*** to give a smaller graph and node set *G*_*s*_ and ***Z***_*s*_, (4) alter *G*_*s*_ (based on *structure* assumption) to create a graph 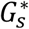 with improper edge connections, (5) calculate 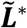 or 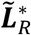 from 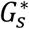, and (5) test for an association between 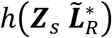 and ***Y*** in the model 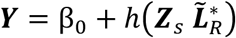. See Figure 4 for a flowchart of these simulation scenarios.

#### Simulated Data

To evaluate PaIRKAT’s overall testing performance and robustness to incorrect pathway information, we simulate data and tests assuming various different types of misspecified pathways. All simulations were accomplished using R(41). Random graphs were generated using the *igraph*(42) package according to the Barabasi-Albert model(43) with *p* nodes representing *p* metabolites within a pathway. The graph’s adjacency matrix was converted into a positive definite precision matrix, **Ω**, using an approached developed by Danaher, et al.(44) and also applied by Shaddox, et al(45). An *n* by *p* matrix of metabolite abundances, ***Z***, was then simulated from a multivariate normal distribution with mean 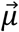 and covariance **Ω**^**−1**^, where 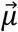 is a vector of *p* 0s. In this way, metabolite connectivity is captured by **Ω**. A continuous outcome *Y*_*i*_ was then simulated from a normal distribution with mean 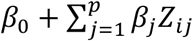 and variance *σ*^2^, where *β*_0_ = 0.2644 and *σ*^2^ = 1.3688^2^. These values for *β*_0_ and *σ*^2^ are drawn from observed metabolomics data. The regularization parameter *τ* is set to 1 for all simulations. All *β*_*j*_ were set to 0 to assess Type I error rates or set to 0.1 to assess power for the twelve different pathway information scenarios described below. Each used 1000 simulations of graphs of size *p* = 15, 30,45 assuming a sample size of *n* = 160, and a testing level of *α* = 0.05 was used for all simulations.

We also conducted a simulation study to assess the impacts of increasing the overall signal to noise ratio on the method’s power. The signal to noise ratio was calculated as the ratio between the overall variance in the mean component, 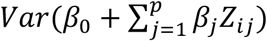, and the overall residual variance, 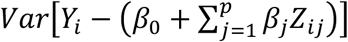. Here *β*_*j*_ is drawn from a normal distribution with a mean and standard deviation of 0.1. Then from normal distributions with gradually increasing means to increase the signal to noise ratio. For simplicity, we only assumed perfect pathway knowledge and simulated under each of the 4 different pathway structures.

### COPDGene Data

We analyzed data collected from the COPDGene study (22), a multicenter observational study that collected genetic data as well as multiple measures of lung function to study chronic obstructive pulmonary disease (COPD). Between 2007 and 2011, 10,198 participants with and without chronic obstructive pulmonary disease (COPD) enrolled (Visit 1). A five-year follow up visit took place between 2013 and 2017 (Visit 2). Blood samples were also obtained for -omics analyses from participants who provided consent. In total, 1136 subjects (1040 non-Hispanic white, 96 African American) participated in a metabolomics ancillary study in which they provide fresh frozen plasma collected using an 8.5 mL p100 tube (Becton Dickson) at Visit 2.

### Metabolomics and Data Processing

P100 plasma was profiled using the Metabolon (Durham, NC, USA) Global Metabolomics platform. Briefly, untargeted liquid chromatography–tandem mass spectrometry (LC–MS/MS) was used to quantify 1392 metabolites and described in(46,47). A data normalization step was performed to correct variation resulting from instrument inter-day tuning differences: metabolite intensities were divided by the metabolite run day median, then multiplied by the overall metabolite median. It was determined that no further normalization was necessary based on the reduction in the significance of association between the top PCs and sample run day after normalization. Subjects with aggregate metabolite median *z*-scores greater than 3.5 standard deviation from the mean (*n* = 6) of the cohort were removed. Metabolites were excluded if >20% of samples were missing values(48). For the 995 remaining metabolites, missing values were imputed across metabolites with k-nearest neighbors imputation (*k* = 10) using the R package *impute*(49). As a final step, metabolomic data was natural log transformed and standardized. Linear regression models were fit to each metabolite controlling for white blood cell count, percent eosinophil, percent lymphocytes, percent monocytes, percent neutrophils, and hemoglobin. The partial residuals were then used as the observed metabolomics data. These data are available at Metabolomics Workbench with identifier PR000907.

Four hundred and thirty six of these metabolites had an id in the KEGG database of human pathways, which was accessed using the *keggLink* function from the *KEGGREST* package(50). These 436 metabolites appear in 161 KEGG pathways, and 28 of these 161 KEGG pathways contained 10 or more metabolites. Edges in a pathway’s graph were defined by connections within a pathway from the KEGG reaction database. Note that our filtered dataset did not contain every metabolite within the 28 KEGG pathways selected, and therefore some of the analyzed pathways have less that 10 metabolites.

### Clinical Variables

We focus on two COPD phenotypes: (1) percent emphysema and (2) the ratio of post-bronchodilator forced expiratory volume at one second divided by forced vital capacity (FEV_1_/FVC). Emphysema, a measure of erosion of the distal airspaces, has been linked with the clinical severity of COPD(51). It is an imaging-based phenotype defined as the 15th percentile lung voxel density in Hounsfield units adjusted for total lung capacity from quantitative CT imaging analyses. FEV_1_/FVC is a measure of airflow obstruction. To normalize FEV_1_/FVC, we use the following log ratio transformation, 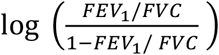. After removing incomplete cases we were left with 1,113 complete cases for the FEV_1_/FVC analysis and 1,065 complete cases for the percent emphysema analysis.

### Analysis

We compared results from tests that included pathway connectivity via 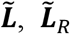, and tests that ignored pathway connectivity for the 28 pathways that had measurements on at least 10 of the metabolites in the pathway. P-values were calculated from a score test as described Section 2 with *τ* = 1. P-values from each method were indistinguishable from one another for both data sets with over 1,000 observations. However, many data sets may not be that large. To demonstrate the differences in performance, 100 random subsets of sizes 100, 200, 300, 400, and 500 were taken from both the log FEV_1_/FVC ratio and the percent emphysema data sets. All three methods were used to test for associations between phenotype and metabolites within a pathway. The 100 p-values were then averaged to measure the performance of each method. All null models included subject age, sex, BMI, smoking status (current, former, never), pack-years of smoking, and the clinical center as covariates.

## Acknowledgements

The project described was supported by Award Number U01 HL089897 and Award Number U01 HL089856 from the National Heart, Lung, and Blood Institute, and U01 CA235488 from the National Cancer Institute. The content is solely the responsibility of the authors and does not necessarily represent the official views of the National Heart, Lung, and Blood Institute, National Cancer Institute, or the National Institutes of Health.

COPDGene is also supported by the COPD Foundation through contributions made to an Industry Advisory Board that has included AstraZeneca, Bayer Pharmaceuticals, Boehringer-Ingelheim, Genentech, GlaxoSmithKline, Novartis, Pfizer, and Sunovion.

