## Supplemental Figures and Tables for "PaIRKAT: A pathway integrated regression-based kernel association test with applications to metabolomics and COPD phenotypes"

### Supplemental Material

Figure S1: Associations between metabolite subsets and log FEV1/FVC ratio. Average p-values from kernel regressing tests that do not include pathway information (No Laplacian, red circles), include pathway information through a normalized Laplacian ($\tilde{\boldsymbol{L}}$, green triangles), and include pathway information through a regularized normalized Laplacian ($\tilde{\boldsymbol{L}}_R = \left(I + \tau\tilde{\boldsymbol{L}}\right)^{-1}$, blue squares) are displayed. P-values were averaged over 100 random subsets of size 100, 200, 300, 400, and 500 from the COPDGene dataset. $\tau$ was set to 1 for all tests that used $\tilde{\boldsymbol{L}}_R$.

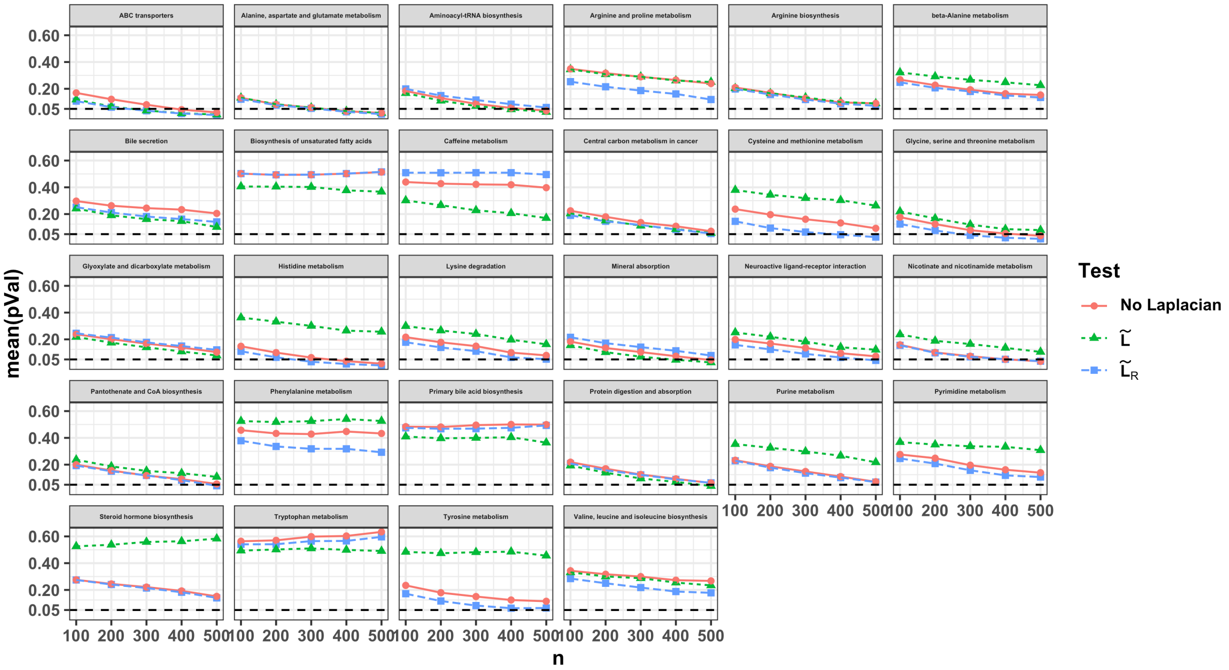

Figure S2: Associations between metabolite subsets and percent emphysema. Average p-values from kernel regressing tests that do not include pathway information (No Laplacian, red circles), include pathway information through a normalized Laplacian ($\tilde{\boldsymbol{L}}$, green triangles), and include pathway information through a regularized normalized Laplacian ($\tilde{\boldsymbol{L}}_R = \left(I + \tau\tilde{\boldsymbol{L}}\right)^{-1}$, blue squares) are displayed. P-values were averaged over 100 random subsets of size 100, 200, 300, 400, and 500 from the COPDGene dataset. $\tau$ was set to 1 for all tests that used $\tilde{\boldsymbol{L}}_R$.

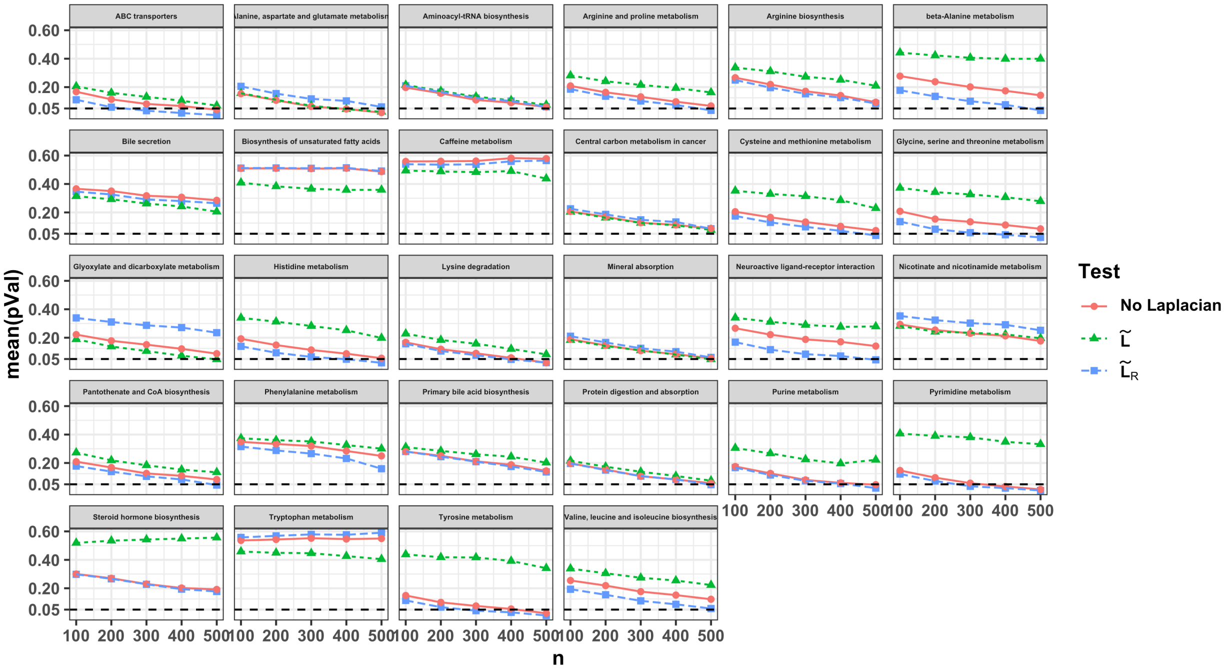


Figure S3: Examples graphs with high, medium, and low edge densities. Low density graphs were generated according the Barabasi-Albert model for graph simulation. Medium- and high-density graphs were generated by giving each unconnected node either a 5% or 15% chance of becoming connected, respectively.

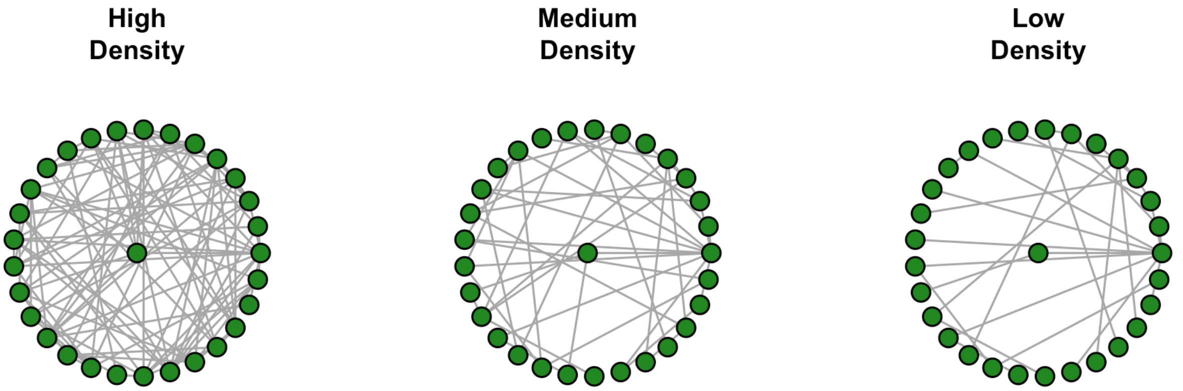

Figure S4: Example of a graph with missing nodes. Graphs were generated according to the Barabasi-Albert model. Then any node with degree below the 25th percentile of degrees within the graph had a 25% chance of being dropped.

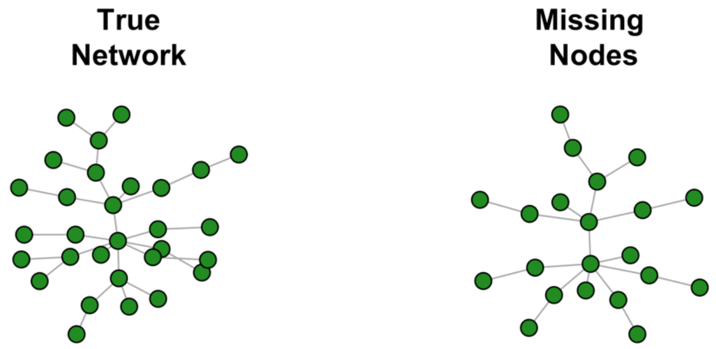

*Complete Pathway*

Table S1: Type 1 error rates using complete pathway. Error rates were calculated from score tests on 1000 simulated data sets. All simulations used graphs with 15, 30, or 45 nodes. No nodes or edges were dropped for these simulations. Pathway information was included in kernel score test through the normalized Laplacian $\tilde{\boldsymbol{L}}$.

| | **Pathway size** | | |
|---|---|---|---|
| | 15 | 30 | 45 |
| *Perfect* | 0.061 | 0.052 | 0.049 |
| *Partial 10%* | 0.059 | 0.055 | 0.044 |
| *Partial 40%* | 0.062 | 0.054 | 0.055 |
| *Partial 70%* | 0.049 | 0.063 | 0.062 |
| *Complete Mismatch* | 0.064 | 0.065 | 0.052 |

*Missing edges*

Table S2: Type 1 error rates using pathways with 5% missing edges. Error rates were calculated from score tests on 1000 simulated data sets using graphs with 15, 30, or 45 nodes. The graph used to simulate $\boldsymbol{Z}$ and $\boldsymbol{Y}$ was of medium edge density, while the graph used to test was of low density. The low-density graphs are drawn from the Barabasi-Albert model with edge density 0.13, 0.07, and 0.04 for graphs with 15, 30, and 45 nodes, respectively. Medium edge density graphs are created by giving any 2 nodes without a direct edge between them a 5% chance of becoming directly connected. This creates graphs with an average edge density of 0.18, 0.11, and 0.09 for graphs with 15, 30, and 45 nodes, respectively. Pathway information was included in kernel score test through the normalized Laplacian $\tilde{\boldsymbol{L}}$.

| | | **Pathway size** | |
|---|---|---|---|
| | 15 | 30 | 45 |
| *Perfect Network* | 0.056 | 0.051 | 0.041 |
| *Partial 10 Mismatch* | 0.059 | 0.063 | 0.042 |
| *Partial 40 Mismatch* | 0.055 | 0.057 | 0.056 |
| *Partial 70 Mismatch* | 0.049 | 0.051 | 0.040 |
| *Complete Mismatch* | 0.068 | 0.054 | 0.056 |

Table S3: Type 1 error rates using pathways with 15% missing edges. Error rates were calculated from score tests on 1000 simulated data sets using graphs with 15, 30, or 45 nodes. The graph used to simulate $\boldsymbol{Z}$ and $\boldsymbol{Y}$ was of high edge density, while the graph used to test was of low density. The low density graphs are drawn from the Barabasi-Albert model with edge density 0.13, 0.07, and 0.04 for graphs with 15, 30, and 45 nodes, respectively. High edge density graphs are created by giving any 2 nodes without a direct edge between them a 15% chance of becoming directly connected. This creates graphs with an average edge density of 0.26, 0.21, and 0.19 for graphs with 15, 30, and 45 nodes, respectively. Pathway information was included in kernel score test through the normalized Laplacian $\tilde{\boldsymbol{L}}$.

| | Pathway size | | |
|---|---|---|---|
| | 15 | 30 | 45 |
| *Perfect Network* | 0.061 | 0.050 | 0.057 |
| *Partial 10 Mismatch* | 0.050 | 0.059 | 0.043 |
| *Partial 40 Mismatch* | 0.059 | 0.045 | 0.056 |
| *Partial 70 Mismatch* | 0.048 | 0.046 | 0.043 |
| *Complete Mismatch* | 0.063 | 0.052 | 0.039 |

*Missing nodes*

Table S4: Type 1 error rates using pathways with dropped nodes. Error rates were calculated from score tests on 1000 simulated data sets using graphs 15, 30, or 45 nodes initially. The graph used to simulate $\boldsymbol{Z}$ and $\boldsymbol{Y}$ contained all nodes. Nodes with degree below the 25$^{th}$ percentile within a graph had a 25% chance of being dropped before testing. Pathway information was included in kernel score test through the normalized Laplacian $\tilde{\boldsymbol{L}}$.

| | Pathway size | | |
|---|---|---|---|
| | 15 | 30 | 45 |
| *Perfect Network* | 0.052 | 0.054 | 0.047 |
| *Partial 10 Mismatch* | 0.053 | 0.055 | 0.045 |
| *Partial 40 Mismatch* | 0.049 | 0.053 | 0.057 |
| *Partial 70 Mismatch* | 0.045 | 0.049 | 0.052 |
| *Complete Mismatch* | 0.049 | 0.052 | 0.040 |
